# Single cell functional genomics reveals the importance of mitochondria in cell-to-cell variation in proliferation, drug resistance and mutation outcome

**DOI:** 10.1101/346361

**Authors:** Riddhiman Dhar, Alsu M Missarova, Ben Lehner, Lucas B Carey

## Abstract

Mutations frequently have outcomes that differ across individuals, even when these individuals are genetically identical and share a common environment. Moreover, individual microbial and mammalian cells can vary substantially in their proliferation rates, stress tolerance, and drug resistance, with important implications for the treatment of infections and cancer. To investigate the causes of cell-to-cell variation in proliferation, we developed a high-throughput automated microscopy assay and used it to quantify the impact of deleting >1,500 genes in yeast. Mutations affecting mitochondria were particularly variable in their outcome. In both mutant and wild-type cells mitochondria state – but not content – varied substantially across individual cells and predicted cell-to-cell variation in proliferation, mutation outcome, stress tolerance, and resistance to a clinically used anti-fungal drug. These results suggest an important role for cell-to-cell variation in the state of an organelle in single cell phenotypic variation.

## Introduction

Isogenic populations often exhibit considerable phenotypic heterogeneity even in an identical environment. One common phenotypic variation that has been observed in isogenic populations of microbial and mammalian cells, including cancer cells is variation in proliferation rate [1-9]. Phenotypic variations that are often coupled with variation in proliferation rate are the abilities of an individual cell to survive stress and drug treatment [3,4]. In this regard, the existence of “persister” cells in microbial populations is well known and poses a significant challenge for antibiotic treatment [2,10–14]. Similarly, individual cells in tumors have been shown to vary in their ability to survive anticancer drugs and can lead to drug-resistant populations [15–20]. Recent advances in single-cell techniques are revealing the extent of transcriptomic and metabolic differences among isogenic cells [21,22]. The existence of such heterogeneity in gene expression in isogenic microbial and animal populations has been shown – to some extent – to underlie the variable outcome of mutations [23–27]. Incomplete mutation penetrance and variable expressivity is also common in human disease [28–31].

Heterogeneity can arise due to stochastic fluctuations in biological processes taking place inside cells. This can happen due to the small numbers of molecules involved in processes such as transcription [32–34] or during stochastic partitioning of cellular components during cell division [35,36]. Although cell-to-cell variation in the expression level of single genes has been correlated with variation in proliferation rate and stress and drug resistance [3,4,15,24–26,37,38], the true underlying causes of such phenotypic heterogeneity are poorly understood.

To identify genes and cellular processes involved in the generation of phenotypic heterogeneity we set up a high-throughput microscopy assay to quantify proliferation heterogeneity in a yeast population. Using this assay, we quantify the impact of deletion of >1,500 genes on proliferation heterogeneity. We present evidence that the variation in mitochondria state is an important determinant of phenotypic heterogeneity in individual cells. We also show that mitochondria state impacts gene expression and stress and drug resistance in individual cells. Taken together, our work suggests an important role for an organelle in generating phenotypic heterogeneity across individual cells in a homogenous environment.

## Results

### Natural and lab yeast populations show proliferation heterogeneity

To investigate cell-to-cell variation in proliferation rates, we developed a high-throughput automated time-lapse microscopy assay that measures the proliferation rates of thousands of single-cells per plate as they grow into micro-colonies. The assay uses a microscope with laser-based autofocus for image acquisition and a liquid handling robot to minimize density-dependent effects on proliferation. The data obtained are highly reproducible with mode proliferation rate of a lab strain being 0.407±0.011 h^−1^, (mean±sd) during >2 years of data collection (n=44 batches; Fig 1A).

**Figure 1:**
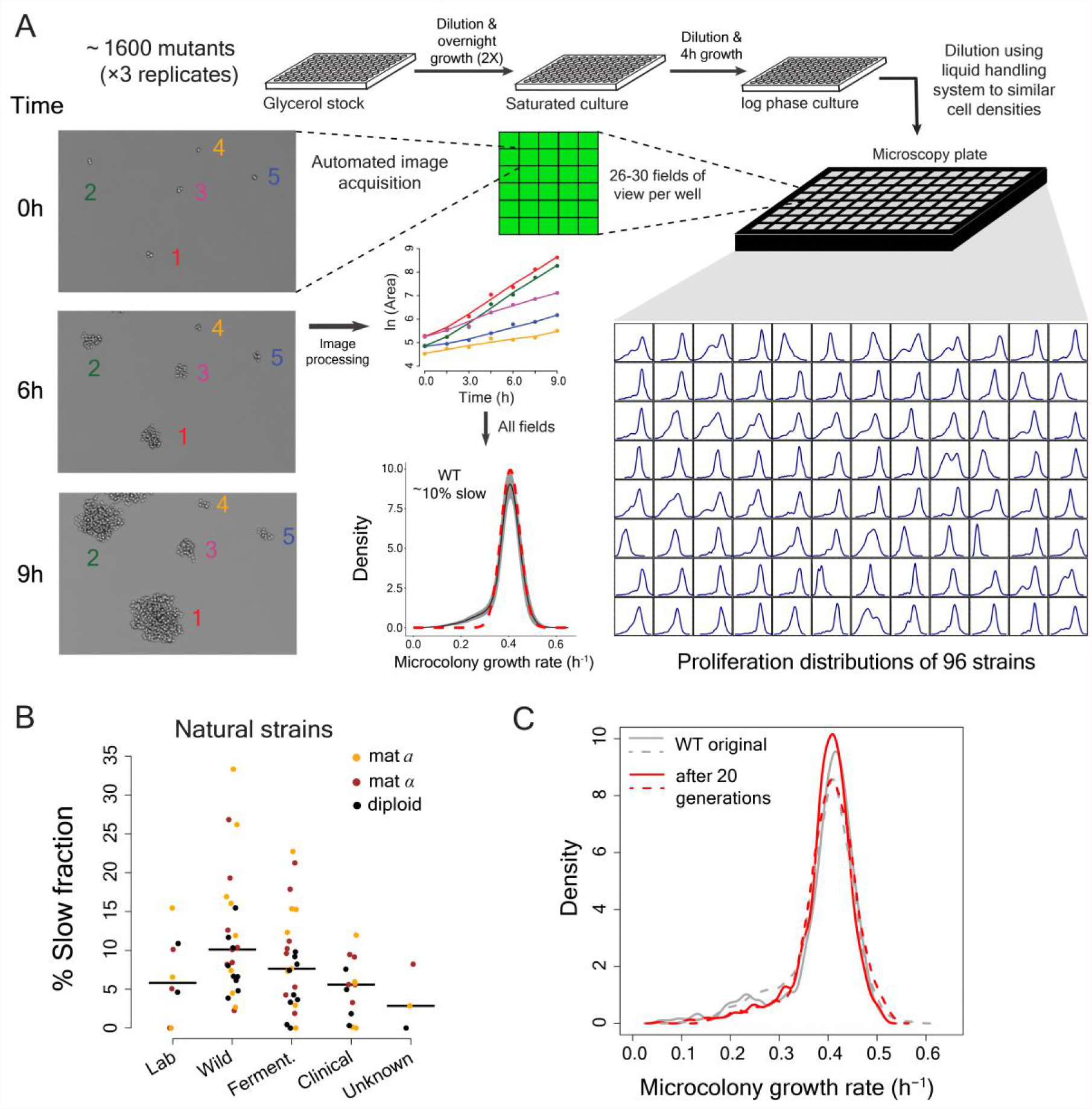
High-throughput analysis of single cell proliferation rate heterogeneity. (A) High throughput microscopy setup – log phase yeast cells were diluted onto conA coated microscopy plate using Biomek NX liquid handling system to have similar cell density across wells. Cells were observed using an ImageXpress Micro system. Images were processed using custom scripts and data for area of microcolony vs. time were obtained. The points in the area vs. time graph show actual data and the solid lines show lowess fits. Data collected from all fields of view in a well constitute a microcolony proliferation rate distribution for a strain. The common lab yeast strain BY4741 (WT) has ~10% slow proliferating sub-population. The density shows mean density and the shaded areas in grey represent ±1 s.d. value at each point. The dotted red line shows the expected proliferation distribution if it were normally distributed. (B) Natural strains of yeast [39] also have slow proliferating sub-populations. Each point represents data for one strain. Solid lines show median value. (C) WT strain re-created the original proliferation distribution even after 20 generations of growth. The plot shows data from two replicate measurements.

Laboratory strains of the budding yeast *Saccharomyces cerevisiae* showed substantial cell-to-cell variation in proliferation, with ~10% of cells forming a slow growing sub-population in defined growth medium (Fig. 1A) [3,39]. This slow growing sub-fraction is not unique to laboratory strains but exists in all natural and clinical isolates that we tested (Fig. 1B; Supplementary table 1) [39]. Growth of the culture for an additional 20 generations did not alter the proliferation rate distribution; the mixture of slow and fast proliferating cells is maintained (Fig. 1C). Proliferation is therefore a stable heterogeneous phenotype within a population, with the amount of heterogeneity depending on the genetic background.

### A genome-scale screen to identify genes that alter proliferation heterogeneity

The effect of individual gene deletions on population-level growth rate has been well studied [40,41]. Many deletions have been shown to reduce population growth rate and can do so in different ways. Deletions can uniformly affect fitness of all the cells or alternatively, can affect fitness of a sub-population whereas the rest of the population remains unaffected. Inter-individual variation in the outcome of mutations has been observed before in multicellular organisms [23–25] but its relative occurrence has not been systematically quantified.

We therefore used the automated microscopy assay to quantify proliferation rate heterogeneity in triplicate for 1,600 gene deletion mutants (Supplementary table 2, including 1,150 gene deletions previously reported as affecting growth rates) [40,41]. We obtained reproducible data (where at least two replicate measurements showed good agreement) for 1,520 deletions, with 1,112 of these reducing the population proliferation rate in our experiment (Mann-Whitney U test, FDR<0.1).

Deletion strains with similar population proliferation rates often showed strikingly different degrees of intra-population heterogeneity (Fig, 2A-C). At the single cell level, ~39% of all mutants with a significant reduction in population proliferation rate (1112 mutants) showed significantly higher variation in mutation outcome compared to the WT strain. Among these mutants, ~13% had the same mode growth rate as the WT strain while showing higher variability. However, almost all mutants (1111 or 1112) had a subset of cells proliferating at the same rate as the bulk of the wild-type (WT) population (one sample Wilcoxon rank-sum test for overlap with bulk WT distribution differing from zero, FDR<0.1; Supplementary fig. 1, Fig. 2D). Thus, a highly variable outcome is actually the normal outcome for proliferation rate at the single cell level when a non-essential gene is inactivated (Fig. 2D, Supplementary fig. 2A).

**Figure 2:**
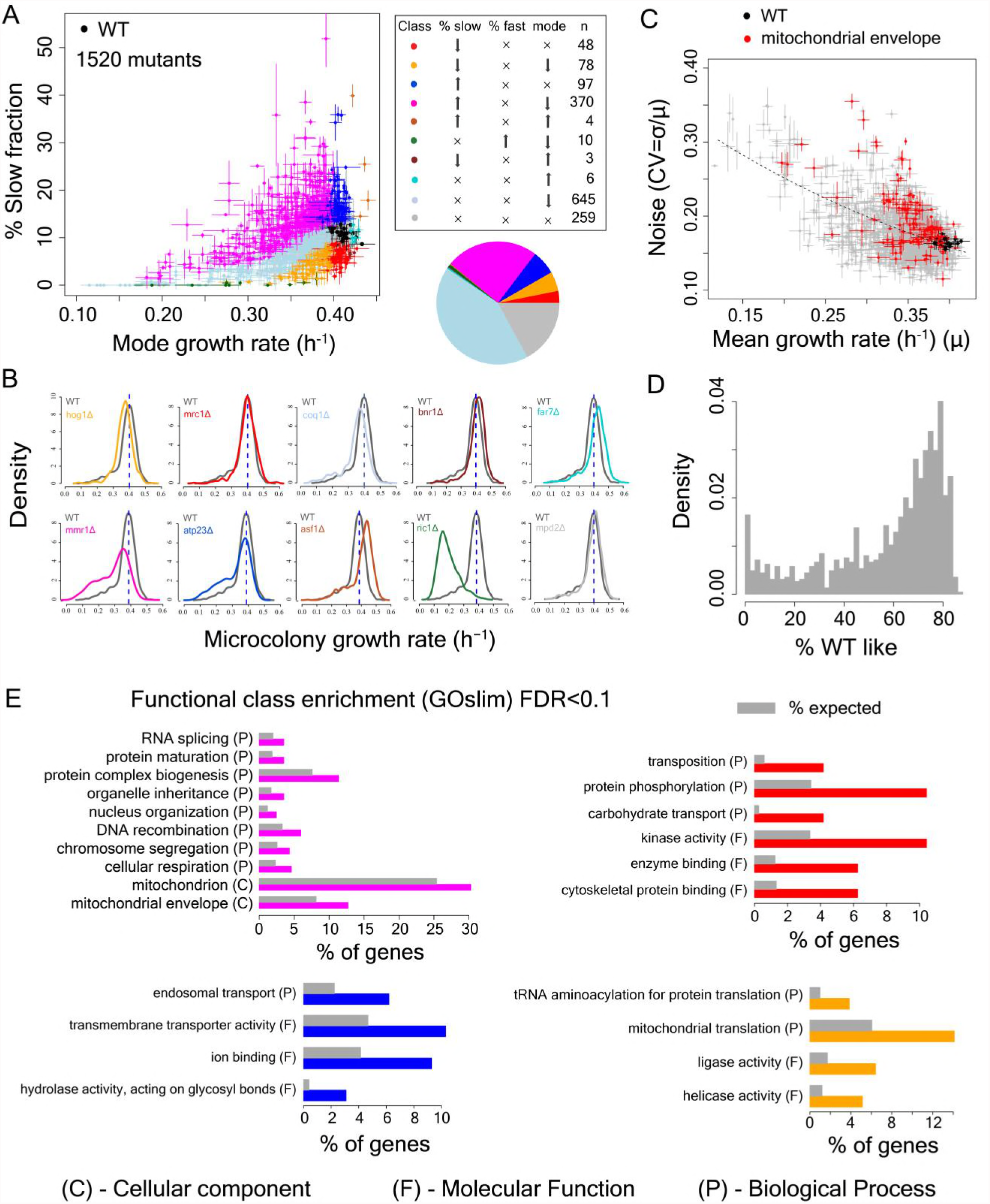
Single cell proliferation rate distributions for 1,500 gene deletions. (A) Mode growth rate (h^−1^) and % slow fraction for 1520 deletion strains. The points represent average values across replicates and the bars represent ± 1 s.d. values. The colours show classification of mutants into different categories according to change in mode growth rate (see Methods, FDR<0.1) and change in % slow fraction (FDR<0.1) compared to the wild-type (WT) strain. The table and pie chart show the number and proportion of strains in each group (colour coded). Replicate data for WT strain are shown by multiple black points. (B) Examples of growth distributions of mutants classified into different groups which are colour coded as in A. The distribution in dark grey shows WT growth distribution. (C) Coefficient of variation (CV) vs. mean growth rate for all strains. WT values are shown in black; mutants of genes that localize to mitochondrial envelope in red. The points represent average values across replicates and the bars represent ± 1 s.d. values. (D) % of WT-like cells in all mutants showing variable mutation outcome. It was calculated for all mutants showing significant reduction in mean proliferation rate and had significant proportion of cells growing as fast as the bulk of the WT proliferation distribution (Wilcoxon rank-sum test). (E) Functional class enrichment (GOslim) analysis for different classification groups show significantly enriched functional classes (Exact binomial test, FDR<0.1). P – Biological Process, F – Molecular Function, C- Cellular Component. Bars show % of genes in a particular group (colour coded) being present in that particular functional class.

### Deletion of genes involved in mitochondrial function alter heterogeneity

To identify the determinants of this cell-to-cell variation in growth-rate and mutational impact we classified each of the deletions by how it affected both the mode and distribution of cellular proliferation rates (Supplementary table 2, Fig. 2A,B). Approximately 17% of the mutants showed no change in either mode proliferation rate or percentage of slow sub-population (in grey), whereas ~43% exhibited a change in mode proliferation rate but no change in slow fraction (in light blue). Interestingly, 48 mutants reduced the slow fraction without any change in mode proliferation rate (in red) and 97 mutants increased the slow fraction without altering the mode proliferation rate (in blue). In addition, there were 78 mutants that reduced both the slow fraction and the mode proliferation rate (in orange). Finally, 370 mutants reduced the mode growth rate but increased the slow sub-population (Fig. 2A). Across mutants, we observed a strong inverse relationship between mean growth rate and noise (co-efficient of variation) (Fig. 2C), as has been observed for gene expression [42,43].

To identify biological processes associated with changes in the slow growing sub-population, we performed a GO functional enrichment analysis on genes in these categories (FDR<0.1). Deletions causing the largest increase in the fraction of slow proliferating cells were highly enriched for nuclear genes encoding mitochondrial proteins (Fig 2C,E). Among the mutants that increased the slow fraction but also reduced mode growth rate (Fig. 2E, magenta), ~30% localized to mitochondria (~1.2-fold enrichment), ~13% localized to the mitochondrial envelope (>1.6-fold enrichment) and ~4.6% were involved in cellular respiration (~2-fold enrichment). In particular, deletion of genes that localized to the mitochondrial envelope resulted in a large increase in slow fraction and noise (Fig. 2E, Supplementary fig. 2B, Supplementary table 2). Mutations that affect mitochondria, and in particular the mitochondria membrane, increase heterogeneity, suggesting that heterogeneity in proliferation might be associated with cell-to-cell variation in mitochondria.

### Mitochondria state but not content predicts slow growth

To further investigate the role of mitochondria in proliferation heterogeneity we used the MitoTracker dye to quantify mitochondrial content in WT cells and five deletion strains with very different proliferation distributions. Total mitochondria content varied little across the strains (Supplementary fig. 2C), ruling out cell-to-cell variation in segregation of the organelle as a driver of heterogeneity. However, signal from the mitochondria membrane potential dye TMRE varied substantially across WT, mutants and natural strains (Fig 3A,B). This suggested that the state of the mitochondria – but not their content – might be driving proliferation heterogeneity.

**Figure 3:**
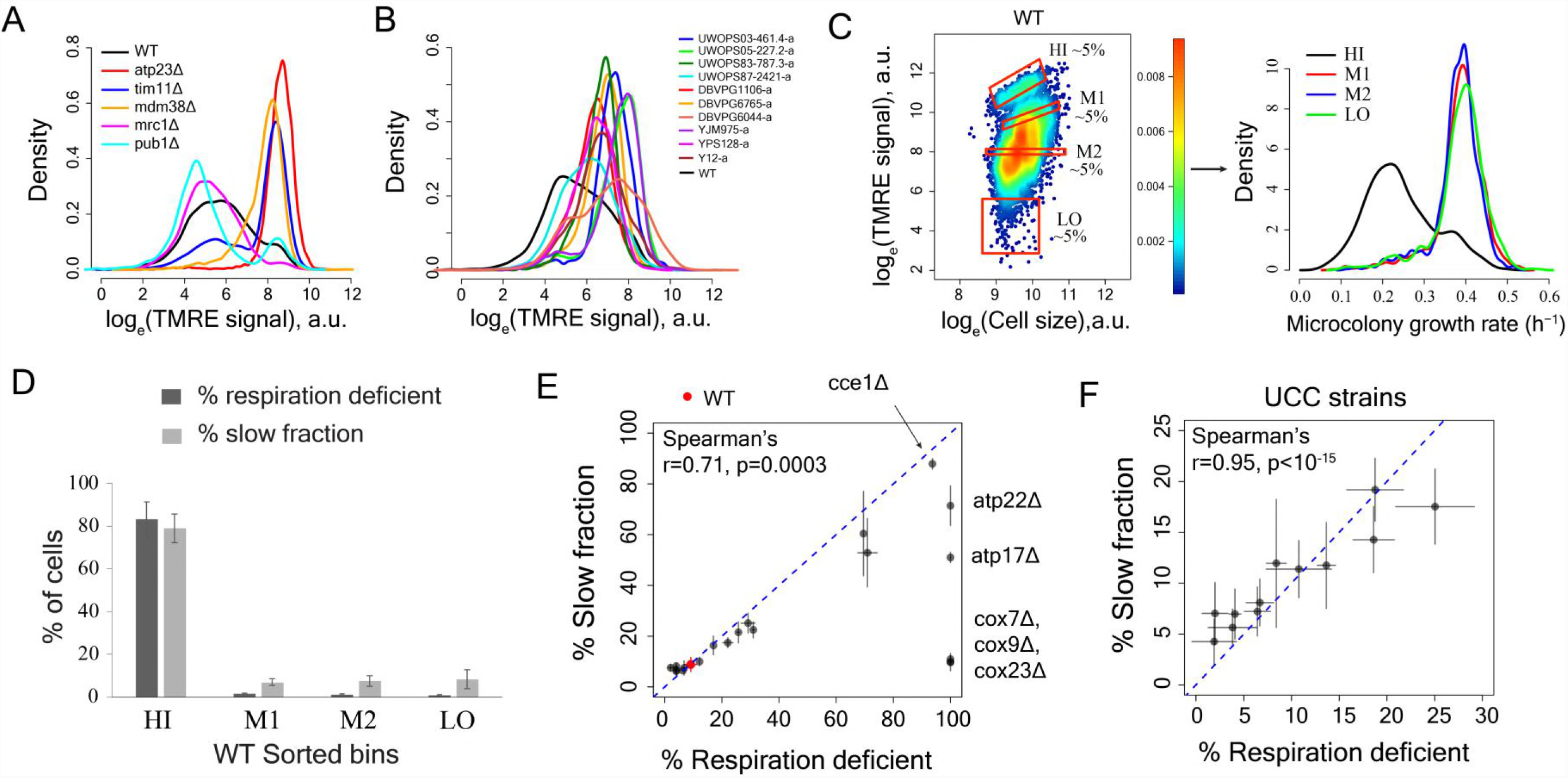
Variation in mitochondria state across single cells underlies proliferation heterogeneity. (A) TMRE stain intensity (log transformed) measured by flow cytometry in WT and deletion mutants. (B) TMRE intensity in WT and natural isolates of *S. cerevisiae* strains. (C) WT cells were sorted by TMRE signal intensity into four bins HI, M1, M2 and LO with gates as shown (~5% of the population sorted in each bin) and growth rate distributions were measured using high throughput microscopy setup. HI bin was enriched for slow growing cells. (D) % of respiration deficient cells in each bin from WT strain. The columns represent the average values from 12 independent experiments and the bars show ±1 s.d. values. (E) Percentage of respiration deficient cells in WT and mutant strains is positively correlated with the percentage of slow growing cells. The blue dotted line represents y=x line. The error bars represent ±1 s.d. measured from at least two biological replicates for each strain. (F) Percentage of respiration deficient cells in UCC strains [44] is strongly positively correlated with the percentage of slow growing cells. The blue dotted line represents y=x line. The error bars represent ±1 s.d. measured from at least two biological replicates for each strain.

To determine if mitochondria state is correlated with variation in growth rate within a population we sorted wild-type cells according to their TMRE staining and measured the fraction of slow proliferating cells. The population with high TMRE was highly enriched for slowly proliferating cells (Fig. 3C). This same fraction was also strongly enriched for respiration deficient cells (Fig. 3D). This strong enrichment for slow proliferation and respiration deficiency was also observed for the cells with high TMRE in gene deletion strains and in natural isolates (Supplementary fig. 3A-D).

We further quantified the TMRE signal and proliferation distribution in a set of twelve strains that differ only by naturally occurring polymorphisms known to affect mitochondrial function by altering mtDNA inheritance [44]. Across all datasets, the percentage of slowly proliferating cells showed a high correlation with the percentage of respiration-deficient cells (Fig. 3E,F) as well as with the percentage of high TMRE cells (Supplementary fig. 4A).

Although cell-to-cell variation in mitochondrial content did not predict proliferation rate variation (Supplementary fig. 4B), mtDNA copy number was substantially lower in the cells with high TMRE (Fig. 4A; Supplementary fig. 5A,B), suggesting a likely role of mtDNA copy number in defining mitochondrial state and ultimately, in generation of growth rate heterogeneity.

**Figure 4:**
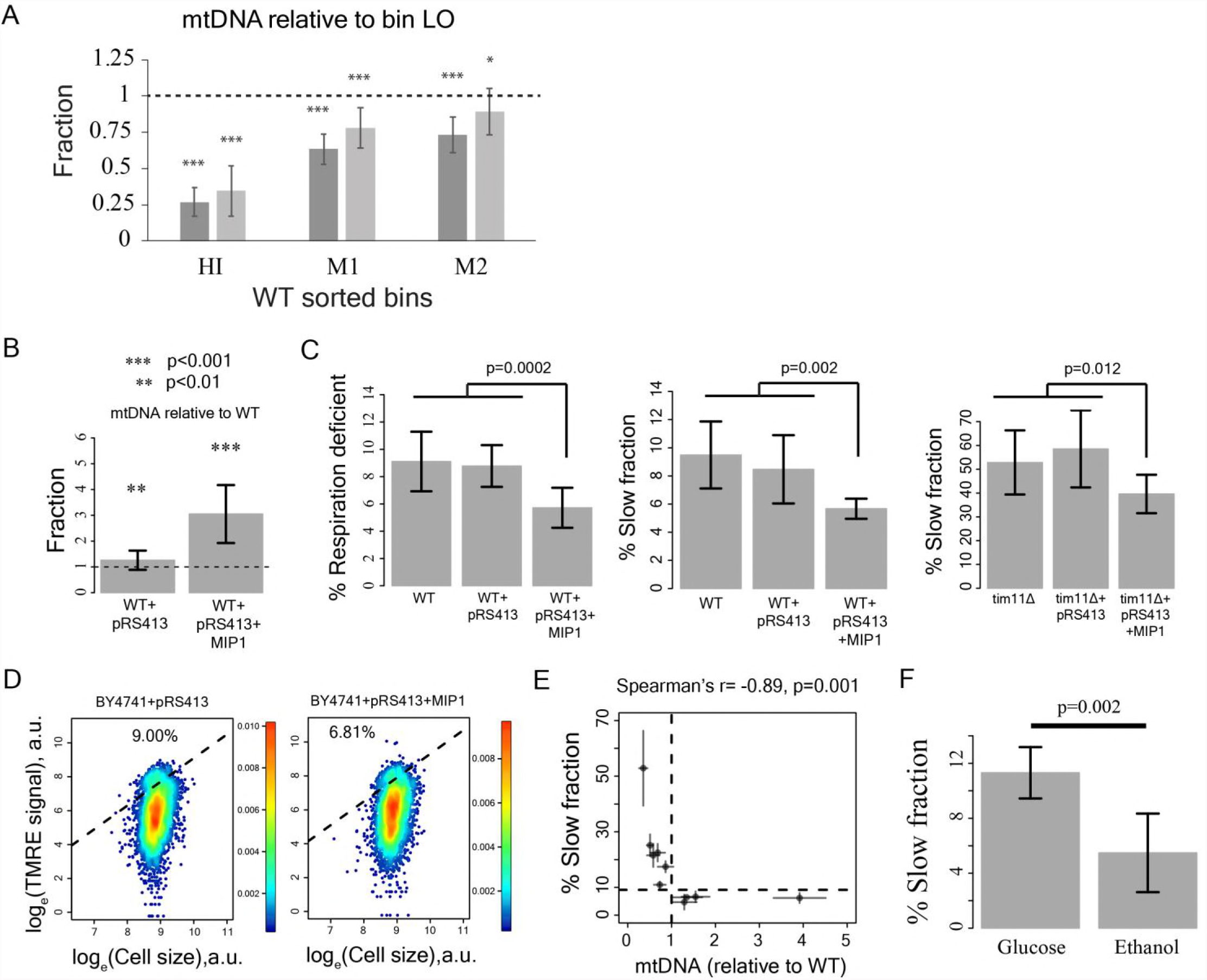
Reduction in mtDNA copy number causes slow growth. (A) mitochondrial DNA (mtDNA) copy number in the sorted bins HI-LO from WT strain measured through quantitative PCR. Two columns show results from two independent experiments. The column represents average mtDNA copy number calculated based on five pairs of primers binding mtDNA and five pairs of primers binding nuDNA and three technical replicates for each of these primers. The bars show ±1 s.d. values. (D) Overexpression of Mip1 gene in WT strain led to significant increase in mtDNA copy number (E) Overexpression of Mip1 gene led to significant reduction in percentage of respiration deficient cells and in slow growing subpopulation in WT strain and reduction in percentage of slowly proliferating cells in tim11Δ mutant. Data are from at least four biological replicates. (F) Overexpression of MIP1 gene in WT strain reduced percentage of cells with high TMRE signal. (G) Percentage of slow growing sub-population was strongly correlated with mtDNA copy number in mutant strains. The dotted lines represent values for WT strain. The error bars represent ±1 s.d. values. (H) Pre-growing WT strain overnight in medium containing ethanol as sole carbon source (that required respiration) reduced percentage of slow growing sub-population by ~50% compared to pre-growth in medium containing glucose as the sole carbon source. Data are from six biological replicates.

To establish a causal relationship between mtDNA copy number and slow growth, we introduced an extra copy of the mitochondrial DNA polymerase Mip1[45], which increased mtDNA copy number 3-fold (Fig. 4B). This reduced both the fraction of slow proliferating and respiration-deficient cells and the fraction of cells with high TMRE signal (Fig. 4C,D), suggesting that variation in mtDNA copy number can be causal for variation in both mitochondria state and proliferation. Consistent with an effect of mtDNA copy number on growth, knocking out of Mip1 gene led to complete loss of mtDNA and resulted in completely slow growing yeast population compared to WT (Supplementary fig. 5C,D) [45]. Furthermore, mtDNA copy number showed a strong correlation with the percentage of slow proliferating cells across mutants (Fig. 4E, Supplementary fig. 5E). Finally, forcing cells to respire by pre-growing them on ethanol as the sole carbon source prior to growth in glucose decreased the fraction of slowly proliferating cells (Fig. 4F). Taken together, these results suggest that alterations in mitochondria state, which can be caused by mtDNA copy number reduction below a threshold and other mechanisms is the underlying cause of slow growth in individual cells.

Cells with high, medium or low TMRE regenerated the original TMRE distribution and growth distribution on 48 hours of growth after sorting (Supplementary fig. 6,7A,B). High TMRE cells could give rise to low TMRE cells and vice versa. Similarly, analysis of growth over a series of time points showed that a small fraction of cells from all sorted bins could switch from the slow growing state to fast growing state and vice versa (Supplementary fig. 7C). However, a subset of cells did not recover their mtDNA copy number or their ability to respire after seven days of growth and remained slow-growing (Supplementary fig. 7D,E). We conclude that cells switch between fast and slow growing states, but that a subset of cells become permanently slow growing, presumably because of an inability to recover functional mitochondria because of mtDNA loss.

### Variation in mitochondrial state predicts additional phenotypic heterogeneity including drug resistance

To systematically understand the physiological differences between sub-populations that vary by mitochondria state, we analyzed the transcriptome by RNA sequencing. Cells with high TMRE have low expression of respiratory and proliferation-associated genes (Fig. 5A, Supplementary fig. 8A,B). Consistent with previous analyses of slow proliferating cells [4,46], they also exhibit a DNA damage response (Fig. 5B) and signs of iron starvation (Fig. 5C), which has previously been reported for respiration deficient cells [47,48]. Cells with intermediate TMRE have very similar proliferation distributions to cells with low potential (Fig. 5C). However, their gene expression was substantially different (Supplementary fig. 8C,D), including reduced expression of respiratory genes (Supplementary fig. 8D).

**Figure 5:**
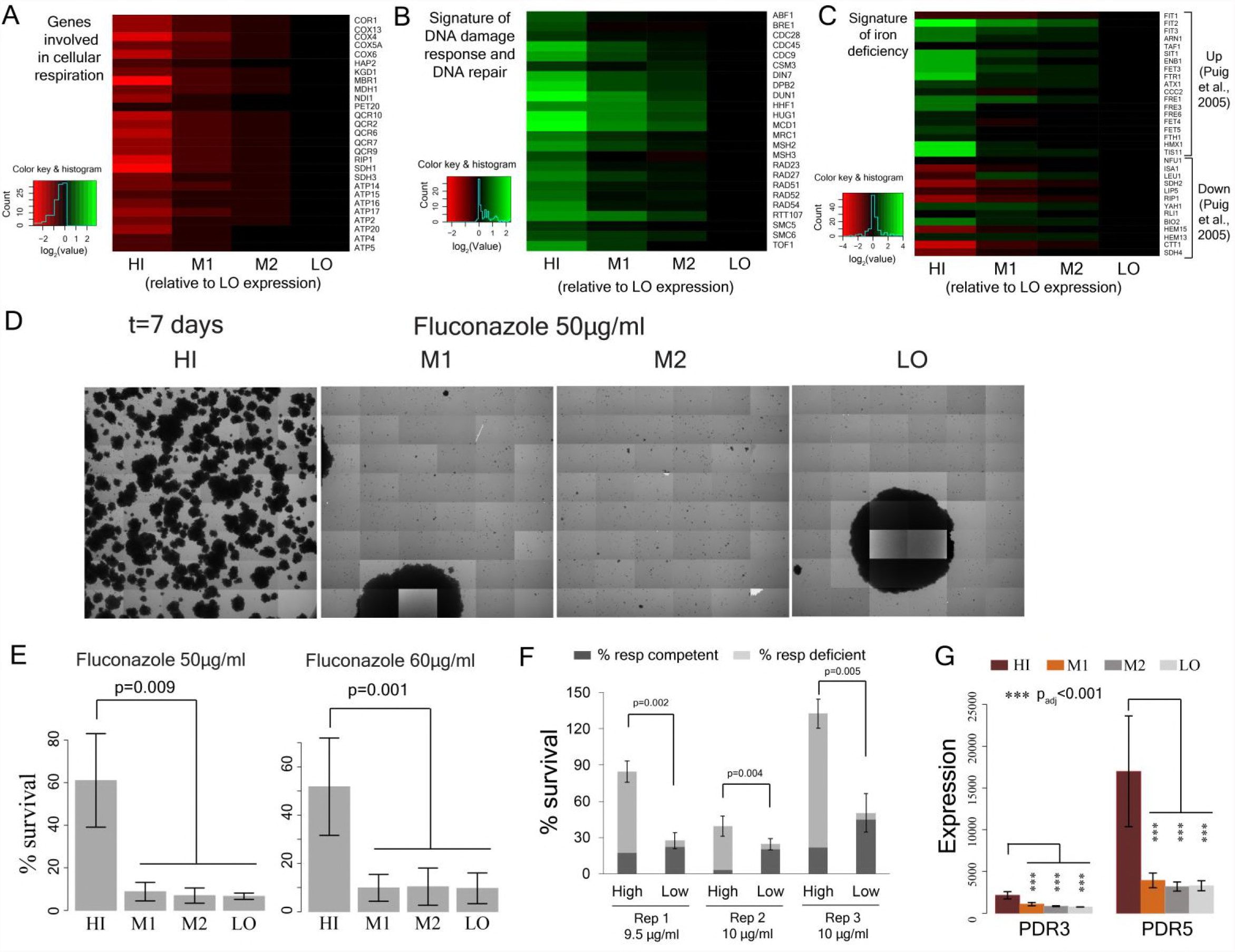
Cell-to-cell variation in mitochondria state predicts single cell drug resistance. (A) Heatmap of expression of respiration genes in cells sorted by their TMRE signal intensity (bins HI-LO). (B) Heatmap of expression of DNA damage response and DNA repair genes. (C) Heatmap of expression of genes associated with iron deficiency [48]. Data are from four independent experiments. (D) Sorted bins from WT cells were subjected to a commonly used antifungal drug fluconazole and were observed under microscope for growth over 7 days. The images show growth of cells in bins HI, M1, M2 and LO in 50μg/ml of fluconazole after 7 days. (E) Cells of HI bin showed significantly higher survival compared to other bins in both 50μg/ml (3 independent experiments) and 60μg/ml fluconazole (4 independent experiments). Cells were grown in liquid medium supplemented with Fluconazole on microscopy plates and viability was calculated from microscopic observations over 7 days. Error bars show ±1s.d. values. (F) Percentage survival of high and low TMRE cells on fluconazole plates. High TMRE cells showed higher survival than low TMRE cells (Mann-Whitney U test). A substantial fraction of surviving high TMRE cells were respiration competent. The error bars represent ±1 s.d. values from 6 technical replicates for each bin. X-axis shows fluconazole concentrations used from 3 independent experiments. (G) From RNA sequencing data, cells from HI bin showed significantly higher expression of multidrug transporter *PDR5* gene and its transcriptional activator *PDR3* compared to cells from bins M1, M2 and LO. Results are from 4 independent experiments.

In bacteria [49] and yeast [3,50,51], slow growing cells can have increased stress resistance. We therefore tested whether the cells with high TMRE in a population are more resistant to acute heat stress. However, cells with high TMRE were more sensitive to heat shock as well as to oxidative stress (Supplementary fig. 9A, B) and expressed some stress-response genes at lower levels (Supplementary fig. 9C).

Slow growing microbes and cancer cells often have increased drug resistance [2,4,12,17,52,53]. Moreover, in several species of fungi, complete loss of mitochondria is associated with elevated resistance to some antifungal drugs [54–56] and respiration deficient strains are often isolated from drug-treated patients [57–59].

We tested therefore whether the high TMRE cells differed in their sensitivity to a clinically used antifungal drug, fluconazole. We found that cells with high TMRE were ~5-7-fold more resistant to high concentrations of fluconazole (Fig. 5D,E; Supplementary fig. 10A). The cells surviving fluconazole treatment included a sub-fraction able to respire (Fig. 5F). Cell-to-cell variation in mitochondria state is therefore also an important predictor of cell-to-cell variation in drug resistance.

Fluconazole targets the cytochrome P450 14α-sterol demethylase enzyme (ERG11), resulting in depletion of ergosterol, a key component of the yeast cell membrane [60,61]. Resistance to fluconazole has been previously reported to depend on the multidrug transporter PDR5 [62–64]. High TMRE cells had significantly higher level of PDR5 expression (Fig. 5G; Supplementary fig. 10B) as well as higher expression of the ergosterol biosynthesis pathway (Supplementary fig. 10D). Consistent with previous work [65,66], the elevated expression of PDR5 in the high membrane potential cells was dependent on the PDR3 transcription factor (Supplementary fig. 10C). Thus, the increased resistance to fluconazole of high TMRE cells is likely to be mediated, at least in part, by increased expression of a multidrug transporter.

## Discussion

In summary, we have shown here that mitochondria state – but not content – varies substantially and reversibly across individual yeast cells and that this is associated with cell-to-cell heterogeneity in proliferation, mutation outcome, and stress and drug resistance. Laboratory strains of yeast have long been known to generate respiratory deficient ‘petite’ colonies at quite high frequency [44,67,68]. However, a slow growing sub-population of cells was observed in all the laboratory, natural, and clinical strains that we tested (Fig. 1A,B). We propose that ‘petite’ colonies are an irreversible extreme phenotype that is generated as part of the more general variation in mitochondria state across single cells that we have identified here.

Although, mitochondrial genes showed the strongest enrichment for an increased slow fraction in our gene deletion screen, other causes of slow growth will, of course, also exist. For example, deletions of genes associated with chromosome segregation and nucleus organization also affected heterogeneity but had no apparent relation to mitochondrial function.

Previous theoretical studies have proposed that variability in the partitioning of cellular components could lead to heterogeneity [35,36]. However, prior experimental work on the fidelity of mitochondria inheritance has shown it to be high, suggesting it is likely to be of little phenotypic consequence for single cells [69]. In contrast, we have shown here that cell-to-cell variation in the *state* of the organelle can be high and predicts phenotypic variation among single cells. Variation in mitochondria state was related to variation in mtDNA copy number in individual cells, but we do not currently know if this is the only – or even the most common – cause of variation in the state of the organelle across single cells. Future work will be required to track down the upstream, proximal causes of this cell-to-cell variation in organelle functional state. The list of gene deletions that alter growth heterogeneity that we have reported here provide a rich resource for this future work. However, it is clear from the results of our screen that many different perturbations to mitochondria – and in particular to the mitochondria envelope and the respiratory complexes – increase this heterogeneity.

Prior work on the causes of variation in proliferation rates, stress and drug resistance, and mutation outcome across individuals and individual cells has focused on fluctuations in gene expression as causative influences [3,4,15,24–26,37,38]. Here we have shown that, in yeast, variation in an organelle is strongly associated with heterogeneity in gene expression across single cells. In animals, inherited and somatic genetic variation in the mitochondrial genome can act as an important modifier of phenotypic variation [70–73]. Recent work has also revealed substantial variation in mtDNA copy number across human tumors [74]. Moreover, in mammalian cells, mitochondrial variability has been suggested to be an important influence on cell-to-cell variation in gene expression and splicing [75–77] and to influence variability in cell death by modulating apoptotic gene expression [20]. Taken together, these results suggest important roles for cellular organelles, in general, and mitochondria, in particular, in the generation of heterogeneity among individual cells. In future work, therefore, it will be important to test the extent to which cell-to-cell variation in the state of mitochondria and other organelles also contributes to variable phenotypic outcomes, mutation effects, and drug resistance in human cells, including in cancer.

## Methods

### High-throughput microscopy assay

96 strains were grown from glycerol stocks in a 96-well plate containing Synthetic Complete medium (0.67% Yeast Nitrogen Base without amino acids and 0.079% Complete Synthetic Supplement (ForMedium, UK)) with 2% glucose (SCD) for 24 hours at 30°C. The cells were diluted 1:50 in fresh medium, grown for 20 hours and diluted again 1:50 in fresh medium. Finally, cells were grown for 4 hours, cell densities were determined by OD at 600nm in a Tecan plate reader and then were diluted to another plate containing SCD or appropriate medium required for microscopy experiment using a Biomek NX (Beckman Coulter) liquid handling robot, capable of pipetting variable volumes of cells across wells in a 96 plate, to a target density of ~17000 cells/μl. This minimized any possible bias due to variability in cell densities among strains. A final 5-fold dilution was done by pipetting 80μl cells onto a pre-coated 96-well microscopy plate containing 320μl of SCD. The microscopy plate was then sealed with LightCycler 480 sealing foils (Roche), cells were spun at 450 rpm for 2 minutes and taken for microscopy observations.

Microscopy plates (96-well glass bottom, MGB096-1-2-LG-L, Brooks Life Science Systems) were coated with 200μl sterile solution of 200μg/ml concanavalin A (type IV, Sigma) at 37°C for 16-18 hours. The solutions were then pipetted out and the plates were washed twice with sterile milli-q water. Plates were dried at 4°C for at least 24 hours. Imaging was performed using an ImageXpress Micro (Molecular Devices) microscope, with laser autofocusing, at an interval of 90 minutes for up to 12 hours. The microscope chamber was maintained at 30°C.

### Image processing

Images were processed using custom scripts written in perl. Yeast cells were identified by juxtaposition of bright and dark pixels (10,36). A pixel was considered ‘bright’ if its intensity exceeded mean+2.2 s.d. value and a pixel was considered ‘dark’ if its intensity was below mean-2.2 s.d. value. In addition, Sobel’s edge detection algorithm [78] was applied for identifying yeast cell boundaries with sharp changes in pixel intensity. Clustering was used to identify the microcolonies and the centroid position for each microcolony was calculated. Microcolonies were tracked through centroid tracking over time. Sudden increase or decrease in centroid number in a time series indicated a failure in image acquisition or image processing and such images were discarded from the analysis. To differentiate cells from cellular debris, residuals of concanavalin A coating etc., two filtering steps were used. First, only objects that were bigger than 50 pixels at the start of observation were considered. Second, the object had to increase its size to greater than 2-fold at the end of observation. Whether neighbouring colonies touch each other at any point in time during microscopy observation was also checked. If they did, they were tracked only up to the time they touched each other. To calculate growth rate, linear regression on natural log-transformed area vs. time for three consecutive time points was performed. This was repeated using a three-point moving window over all time points. Of all these regressions, the maximum value was chosen as the microcolony growth rate to avoid biases because of slow down of growth during lag and/or due to possible substrate limitation near the end of observation.

### Screening of deletion mutants, classification and functional enrichment analysis

Growth distributions for deletion mutants were measured in three independent experiments on different days. To calculate reproducibility of growth rate between replicates, mean growth rate was allowed to vary up to 0.05 h^−1^ and then Kolmogorov-Smirnov distance (K-S distance) [79] was calculated between all replicates after shifting one of them (through addition/subtraction) by difference in mean growth rates. Three replicates were considered as three nodes in a graph with K-S distance between them as the edge weight. If the K-S distance between two replicates exceeded 0.1, no edge was drawn between those two nodes. The sub-graph where the maximum number of nodes was connected to each other directly and via shortest possible distance was considered as reproducible replicates. Only mutants with at least two reproducible replicates were considered in our analysis. The number of reproducible replicates for each mutant is given in supplementary table 2.

To calculate slow fraction from a proliferation distribution, first, a cumulative distribution function (cdf) was calculated with density being calculated at an interval of 0.01 h^−1^. The cdf function was then scanned for maximum slope using a window of 5 points. At the point with maximum slope, a line with the maximum slope was fitted and the points that deviated from the fitted line by >0.02 h^−1^ were considered as the edges of the main sub-population. In the next step, if the left sub-population was bigger than the right sub-population, the percentage of slow fraction was calculated as (% left sub-population-% right sub-population) and the percentage of fast sub-population was set to zero. If the right sub-population was bigger than the left sub-population, the percentage of fast fraction was calculated as (% right sub-population-% left sub-population) and the percentage of slow sub-population was set to zero.

If the mode of a growth distribution is reduced (compared to WT) and the growth rate of slow fraction is not reduced, the main sub-population growth distribution is likely to overlap with and mask a slow growing sub-population. To avoid such scenarios, all the reproducible growth distributions for the WT strain were collected and the mode growth rate was computationally reduced in steps of 0.01 h^−1^ without changing the growth rate of the slow sub-population. The percentage slow fraction was calculated at each step. As expected, reduction in mode growth rate without moving the slow fraction led to a reduction in % of slow fraction (Supplementary fig. 2A).

Mutants with altered mode proliferation rate compared to WT strain were identified through Mann-Whitney U test (FDR<0.1). Mutants with altered slow fraction were identified by Mann-Whitney U test (FDR<0.1) after correcting for any change in mode growth rate (Supplementary fig. 2A). Comparison between replicate measurements of mode growth rate and percentage of slow fraction was done (Supplementary fig. 11A). The mean proliferation rate of mutant strains obtained in our assay was comparable with published values (Supplementary fig. 11B).

We used GOslim gene annotation [80,81] for functional class enrichment analysis and we performed a hypergeometric test as follows. Let us assume that in a group ‘g’ from screening (for example, the group with increased slow fraction but no change in mode growth rate), out of total N_g_ genes, X_g_ genes are associated with function *f* according to GOslim annotation. Let us also assume that out of total N genes screened in our data, X genes belong to the functional class *f* according to GOslim annotation. Thus, the probability that the group ‘g’ contains more number of genes of functional class *f* than expected by chance alone is given by 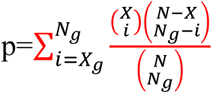, which gives the p-value. A further multiple testing correction was done using Benjamini-Hochberg procedure with FDR<0.1.

### Quantification of incomplete penetrance

Incomplete penetrance was calculated for all mutants that showed significant reduction in mean proliferation rate compared to the WT strain (Mann-Whitney U test, FDR<0.1). For each of these mutants, replicate proliferation distributions were compared with replicate proliferation distributions of WT strain that were reproducible across the screening experiment. Average overlap of the mutant proliferation distributions with the bulk sub-population of each of the WT proliferation distribution was calculated. For WT strain, to calculate bulk sub-population from a proliferation distribution, first, a cumulative distribution function (cdf) was calculated with density being calculated at an interval of 0.01 h^−1^. The cdf function was then scanned for maximum slope using a window of 5 points. At the point with maximum slope, a line with the maximum slope was fitted and the point that deviated from the fitted line by > −0.02 h^−1^ was considered as the edge of the bulk sub-population. Thus, for each mutant, this led to a distribution of a percentage of cells showing WT-like proliferation (Supplementary fig. 1). In the next step, it was tested whether the distribution of percentage of WT-like cells was significantly different from zero (Wilcoxon rank-sum test for one sample) and an FDR correction for multiple testing was performed (FDR<0.1).

### Mitotracker green and TMRE staining

To perform mitotracker green (MitoTracker Green FM, Molecular Probes, Thermo Fisher Scientific) staining, cells were centrifuged at maximum speed for 2 minutes and washed twice with buffer containing 10mM HEPES (pH 7.4) and 5% glucose. Cells were then re-suspended in the same buffer and Mitotracker Green (10μM stock dissolved in DMSO) was added to a final conc. of 100nM. Cells were incubated for 20 mins at 30°C, washed twice with PBS (pH 7.4) and quantified by flow cytometry (LSR Fortessa, BD Biosciences). For TMRE (Molecular Probes, Thermo Fisher Scientific) staining, cells were washed twice with PBS, were re-suspended in PBS, and TMRE was added to a final conc. of 100nM from a 10mM stock dissolved in DMSO. Cells were incubated at 30°C for 30 minutes, were washed twice with PBS and were analysed by flow cytometry or were sorted. There was a gap of 15-20 minutes between the end of staining and beginning of flow cytometry experiments due to the time required for cleaning, priming and setting up of flow cytometry machine parameters. Day-to-day variations were observed in measurement of TMRE distributions.

### Cell sorting and growth measurement of sorted bins

Cells were sorted by TMRE signal into four bins HI, M1, M2, LO (Fig. 3C) in an Aria II SORP cell sorter (BD Biosciences). For growth rate measurement, stress resistance measurement, and mitochondrial DNA quantitation by qPCR in the sorted bins, 100,000 cells per bin were sorted at room temperature into 1.5ml tubes pre-filled with 600μl of PBS. After sorting, 200μl of YPD was added to each tube, cells were centrifuged for 5 mins at maximum speed at room temperature, and the supernatant was thrown away. Cells were re-suspended in 600μl of PBS before proceeding for subsequent experiments. For heat shock experiments, 100μl of sorted cells were put into PCR tubes and were subjected to heat shock in a PCR machine, put on ice for 1 min before measurement of growth and viability. For RNA sequencing experiments, 750,000 cells per bin were sorted in three 1.5ml tubes, each pre-filled with 800μl of PBS. After sorting, 200μl of YPD was added to each tube, centrifuged at maximum speed for 5 minutes, and supernatants were discarded. The cell pellets were gently washed twice using PBS. Total RNA was isolated using MasterPure yeast RNA isolation kit (Epicentre) following manufacturer’s protocol.

To determine percentage of respiration deficient cells, cells were plated on plates containing Synthetic Complete medium with 3% glycerol and 0.1% glucose (SCDG) solidified with 1.5% agar to a target density of ~100-150 colonies per plate. After 5-7 days, number of small and big colonies were counted and the percentage of respiration deficient cells were determined as - percentage of respiration deficient cells 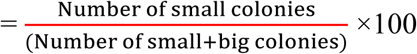

To test for switching of microcolony growth from slow to fast state or from fast to slow state, microcolony growth rates were calculated for each time point using linear regression with data from 3 time-points and with R^2^>=0.9. Microcolonies for which growth rate changed from over 0.3 h^−1^ to less than 0.3 h^−1^ over time were counted as microcolonies switching from fast to slow state. For fast to slow state transition, the transition must happen before last 4 time-points so as to discount slowdown due to nutrient deprivation. Microcolonies for which growth rate changed from less than 0.3 h^−1^ to more than 0.3 h^−1^ over time were counted as microcolonies switching from slow to fast state. For this switching, the transition must happen after first 4 time-points so as not to count lag to log phase switching.

### Measurement of mtDNA copy number by qPCR

To determine mtDNA copy number per cell using quantitative PCR (qPCR), five primer pairs specific to nuclear DNA (ACT1, ALG9, KRE11, TAF10, COX9) and five primer pairs specific to mitochondrial DNA (COX1, ATP6, COX3, ATP9, tRNA – primer picked around tQ(UUG)Q gene) were used (Supplementary table 3). A standard curve for each of primer was made, using six concentrations of genomic DNA serially diluted from the highest concentration by 4-fold at each step. Absolute quantification of DNA copy number was performed using the standard curve. Three technical replicates for each primer and for each sample were set up totaling 30 reactions per sample. To compare mtDNA copy number across sorted bins, nuclear DNA and mtDNA copy numbers in all bins were normalized by the respective values for LO bin. Two sample t-test was used to check whether the normalized value for nuclear DNA differs significantly from the normalized value for mtDNA and a p-value was calculated using a two-sample t-test. Mean mtDNA copy number per cell was calculated by the ratio of mtDNA to nuclear DNA and standard deviations were calculated by taking error propagation models into account.

To overexpress the MIP1 gene, the MIP1 gene under the control of the native promoter (930bp upstream and 262bp downstream, total insert length - 4957bp) was cloned into pRS413 plasmid and then transformed into NEB 10B electrocompetent *E. coli* cells. The plasmid with the verified construct was then isolated and transformed into yeast cells.

### RNA sequencing experiment and data analysis

Isolated total RNA (using MasterPure yeast RNA isolation kit (Epicentre)) was checked and quantified using bioanalyzer. 200ng of total RNA for each sample was taken and was mixed with 4μl of 1:1000 dilution of ERCC spike-in mix1 (Thermo Fisher Scientific). Sequencing was done in Illumina HiSeq with paired end 2×50bp reads. Quality of the sequenced reads was checked using FastQC [82] and then the reads were mapped to reference yeast transcriptome (R64-1-1 reference cdna sequence from Ensembl [83]) using bowtie2 [84]. Mapping statistics was calculated using a custom script where only read pairs mapping concordantly and uniquely to the reference sequence were considered. The data were normalized using ERCC spike-in reads as controls using RUVg method in R package RUVSeq [85]. Correlation between replicates were checked through distance heatmap and PCA analysis (Supplementary fig. 12A,B), using R package DESeq2 [86]. Differentially expressed genes were identified using package DESeq2. Functional enrichment analysis on sets of differentially expressed genes was done using a hypergeometric test as described above with multiple testing correction (FDR<0.1) (Benjamini-Hochberg method) with GOslim gene annotations.

### Reconstruction of single mutants

Gene deletion mutants were remade in the WT strain using sequence specific homologous recombination. First, the deletion cassette from the appropriate deletion strain from the collection was amplified using primers such that the amplified region contained the deletion cassette with KanMX marker and 50-300bp of overhang on either side of the cassette. Particular care was taken to avoid neighbouring genes from being amplified. The PCR product was transformed into competent yeast cells (prepared using lithium acetate and PLI – made by mixing 1ml water, 1ml 1M lithium acetate and 8ml 50% PEG3350) and colonies were selected on G418 plates. Two verified clones for each mutant were picked for experiments. Some of the mutants associated with mitochondrial function were found to be compensated in the deletion collection (Supplementary fig. 12C,D). Beyond the initial high-throughput measurement of proliferation distribution of deletion mutants, all experiments were performed with freshly made deletion mutants.

### Long-term microscopy-based growth measurements

To observe growth of yeast cells in drug (fluconazole dissolved in DMSO, stock conc. 5mg/ml) over 7 days, yeast cells were imaged under the microscope every ~24 hours. To have a bigger part of a well imaged and to increase the number of data points, 48 fields of view per well were imaged in these experiments. Yeast cells and microcolonies were identified as above. Before tracking the microcolonies over time, the images for all fields of view in a well were merged which allowed tracking of microcolonies even if the plate was positioned slightly differently in the microscope at different time points. Microcolonies were tracked over time and growth rates were calculated. A growth rate of 0.02 h^−1^ after first time point was taken as cut-off for survival on fluconazole, as most colonies showed initial growth but then stopped growing. For calculating % survival in heat shock and hydrogen peroxide treatment, the number of colonies showing growth under stressed condition compared to the total number of colonies showing growth under unstressed condition was considered.

### Measurement of respiration capability in drug resistant cells

To test whether the cells that survive fluconazole treatment can still respire, the drug resistance in the sorted cells from the HI and LO bins were measured on agar plates after 15 days of growth in SCD medium supplemented with fluconazole (9.5 or 10µg/ml) and solidified with 1.5% agar. This assay needed lower concentrations of fluconazole compared to the microscopy-based assay, as only the colonies that divided multiple times were visible on the plate. Sorted cells from bins HI and LO were plated directly after sorting onto the drug plates (5-6 replicates per bin), onto plates without any drug as well as onto SCDG plates to calculate the percentage of cells capable of respiration. Cells were counted after 15 days and 40-50 colonies from each plate were randomly picked and checked for respiration capability by plating onto plates containing 3% glycerol as the carbon source.

## Data availability

RNA-sequencing data that support the findings of this study have been deposited in NCBI GEO with the accession code GSE104343 and reviewer token “mtabqsqebncphaz”.

## Code availability

Custom codes for analyzing microscopy images are available at https://github.com/riddhimandhar/MicroscopyCode.git

## Acknowledgements

Work in the lab of BL was supported by a European Research Council Consolidator grant (616434), the Spanish Ministry of Economy and Competitiveness (BFU2011-26206 and SEV-2012-0208), the AXA Research Fund, the Bettencourt Schueller Foundation, Agència de Gestió d’Ajuts Universitaris i de Recerca (AGAUR), Framework Programme 7 project 4DCellFate (277899), and the European Molecular Biology Laboratory–Center for Genomic Regulation (CRG) Systems Biology Program. Work in the lab of LBC was supported by MINECO (BFU2015-68351-P) and AGAUR (2014 SGR 0974) and the Unidad de Excelencia Maria de Maeztu (MDM-2014-0370). The authors thank Dr. Raul Gomez Riera and the CRG Advanced Light Microscopy Unit for assistance with high-throughput microscopy, the CRG/UPF FACS facility for help with flow cytometry, the CRG Genomics Unit for RNA-seq and DNA sequencing experiments, and the UPF Genomics facility for help with automated liquid handling. RD was partially supported by the Swiss National Science Foundation Early Postdoc Mobility fellowship.

## Author contributions

RD set up high-throughput microcopy assays and developed pipelines for image processing. RD and AMM performed high-throughput microscopic proliferation measurements of the deletion mutants and analyzed the data. RD performed the remaining experiments and analyzed the data. RD, LBC, and BL wrote the manuscript with input from AMM. All the authors read the manuscript and approved it.

---

**Supplementary figure 1** – Schematic diagram showing calculation of %WT like cells from mutant proliferation distributions.

**Supplementary figure 2** – (A) To test how reduction in mode growth rate without any change in growth rate of slow sub-population might influence our capability to detect slow fraction, the main sub-population of WT strain was computationally moved by reducing mode growth rate in steps of 0.01 h^−1^ without altering the growth rate of the slow sub-population. This was done for all independent measurements of growth rate distributions for WT. The brown points show average value for all such independent computations and the error bars show ± 1 s.d. values from all computations. The blue points show the %WT-like cells vs. mode growth rate (h^−1^) for mutants exhibiting incomplete penetrance (IP). A mutant was considered to be incompletely penetrant if its proliferation distribution had significant overlap with the bulk of the WT proliferation distribution. (B) Distribution of Noise (Coefficient of variation or CV) (left) and distribution of percentage slow fraction (right) for knock-out of genes that localize to mitochondrial envelope (in red), genes that localize to mitochondria (excluding mitochondrial envelope) (blue) and rest of the genes (black) in our dataset. P-value in red is from statistical test between red and black distributions and P-value in blue is from statistical test between blue and black distributions (Mann-Whitney U test). (C) Mitochondrial content of WT and mutant strains as measured by Mitotracker green intensity using flow cytometry.

**Supplementary figure 3** – (A) TMRE staining and sorting of BY4743 diploid strain. Bin HI showed higher percentage of slow cells in high throughput microscopy assay. Bin HI showed enrichment for respiration deficient cells. The results are from two independent experiments. (B) Sorting of mutant strains based on TMRE intensity and subsequent growth rate measurements showed slower growth in high TMRE cells (left). Cells with high TMRE signal were also enriched for respiration deficient cells in mutants (right). (C) Cells showing high TMRE intensity in Y12 strain (Mat *a* derivative) were enriched for slow growing fraction and respiration deficient cells. (D) Cells showing high TMRE intensity in YJM975 strain (Mat *a* derivative) were enriched for slow growing fraction and respiration deficient cells.

**Supplementary figure 4 –** (A) Percentage of slowly proliferating sub-population was significantly correlated with percentage of high TMRE cells across WT, deletion mutants and UCC strains. (B) Sorting WT cells by Mitotracker green intensity did not enrich for slow growth or percentage of respiration deficient cells in any sorted bin.

**Supplementary figure 5** - (A) From DNA sequencing, fraction of total reads mapping to mtDNA sequence in HI-LO bins from WT strain. The results are from three independent experiments. (B) mitochondrial DNA (mtDNA) copy number in the sorted bins HI-LO from a natural strain (YJM975 Mat *a*) and UCC8363 strain measured through quantitative PCR. (C-D) Knocking out Mip1 gene encoding for a mitochondrial DNA polymerase led to complete loss of mtDNA and also made the entire yeast population slow growing [45]. (E) PCA analysis of WT and mutant strains with % slow fraction, % high TMRE cells, % respiration deficient cells and mtDNA copy number.

**Supplementary figure 6** – Measurement of TMRE in cells of sorted bins HI-LO from WT strain (A) right after sorting (B) after 24 hours and 48 hours growth in SCD medium. (C) Measurement of mitochondrial content using Mitotracker green in sorted bins HI-LO from WT strain after 24 hours and 48 hours growth in SCD medium.

**Supplementary figure 7** – (A) Percentage of respiration deficient cells in sorted bins HI-LO from WT strain right after sorting (left), following 24 hours of growth in SCD medium (middle) and after 48 hours of growth in SCD medium (right). (B) Proliferation distribution measurement after 48 hours growth of sorted bins HI-LO from WT strain. (C) Cells from HI, M1, M2 and LO bins were tracked for growth rate switching (Slow to Fast and Fast to Slow). Each data point represents result from a replicate experiment. (D) mtDNA copy number (using quantitative PCR) of colonies from sorted bins (HI-LO) from WT strain after 7-day growth on SCD plate. (E) Growth rate distribution measurement of small (respiration deficient) and big (respiration competent) colonies from high TMRE bin of WT strain after 48 hours of growth in SCD. Colonies were picked from SCDG plates after 7 days of growth and each curve represents data from one clone.

**Supplementary figure 8** – (A) Functional classes showing significant enrichment among genes over-expressed in cells of bin HI compared to LO (Exact binomial test, p<0.01) and significantly enriched functional classes among genes showing lower expression in cells of bin HI compared to bin LO and the corresponding fold-change in expression. (B) Heatmap of expression of genes involved in cytoplasmic translation. (C) MA plot [86] showing mean expression of genes vs. fold change in expression for comparison between cells in bin LO and bin M1. Differentially expressed genes are marked in red (FDR<0.1). (D) Functional classes showing significant enrichment among genes over-expressed in cells of bin LO compared to M1 (p<0.01) and significantly enriched functional classes among genes showing lower expression in cells of bin LO compared to bin M1.

**Supplementary figure 9** - (A) Left - % survival of cells from sorted bins HI-LO from WT strain after heat shock at 50°C for 2mins, 3mins and 4mins (Mann-Whitney U test). Right - Growth rate of cells in sorted bins HI, M1, M2 and LO from WT strain in SCD medium without any heat shock, after heat shock at 50°C for 2mins, 3mins and 4mins. (B) % survival of cells from HI-LO bins in 0.6mM, 1mM and 1.2mM hydrogen peroxide (Mann-Whitney U test) (C) Expression of some key heat shock and stress response genes in bins HI-LO from RNA sequencing (data from 4 independent experiments).

**Supplementary figure 10** – (A) Percentage survival of sorted cells from bins HI, M1, M2, and LO from diploid BY4743 strain in 50µg/ml of antifungal drug fluconazole. Cells from bin HI exhibited higher drug survival (results from two independent experiments). (B) PDR5-GFP strain was stained with TMRE and a sub-population of cells with high TMRE signal (corresponding to bin HI) also showed higher expression of *PDR5* gene. (C) Deletion of the transcriptional activator *PDR3* in PDR5-GFP strain wiped out the sub-population of cells showing higher expression of PDR5 gene. (D) Heatmap depicting expression of ergosterol biosynthesis genes in bins HI-LO sorted from WT cells.

**Supplementary figure 11 -** (A) Correlation between replicate measurements of mode growth rate of all strains and between replicate measurements of percentage slow fraction as well as between mode growth rate and percentage slow fraction. The panels below the diagonal show plots with the actual data points and the numbers above the diagonal show correlation values (generated using R package corrgram). (B) Correlation of mean growth rate for deletion mutants from our experiment and published data. The pie charts below the diagonal show magnitude of correlation and the numbers above the diagonal show the actual correlation values.

**Supplementary figure 12 -** (A) Heatmap depicting distance between replicate RNA-seq samples of the sorted bins HI-LO of the WT strain. One replicate of LO bin showed high distance from all clusters. (B) PCA analysis of expression level of all genes for all replicates of bin HI-LO. Again, one replicate of bin LO was an outlier which was discarded before subsequent analyses. (C) Growth distributions for mip1Δ strain from the deletion collection and for two remade clones. The solid and the dotted lines represent replicate measurements on different days. The data clearly shows that the growth rate in the strain from the collection is compensated. (D) Growth distributions for mrpl8Δ strain from the deletion collection and for two remade clones.

**Supplementary table 1 -** Mean and Mode growth rate (h^−1^) and % slow fraction for the natural yeast strains from SGRP collection.

**Supplementary table 2 –** Mean, median and mode growth rates (h^−1^), Standard deviation (SD), Noise (Coefficient of variation, CV), % slow fraction, number of replicates showing reproducible results and the classification colour code (as in Figure 2A) for all the mutants with reproducible results.

**Supplementary table 3 –** Primer pairs used for quantifying mtDNA copy number using quantitative PCR.

**Supplementary information 1 –** Proliferation distributions of 1520 deletion mutants for which reproducible measurements were obtained. Multiple lines in each plot represent reproducible replicate measurements. x-axis represents microcolony growth rate (h^−1^) and y-axis represents density.

**Supplementary information 2 –** An example of gating strategy used for cell sorting experiments.

